# Deciphering the coupling state-dependent transcription termination in the *Escherichia coli* galactose operon

**DOI:** 10.1101/2024.08.28.610070

**Authors:** Monford Paul Abishek N, Xun Wang, Heung Jin Jeon, Heon M. Lim

## Abstract

The distance between the ribosome and the RNA polymerase active center, often referred to as the mRNA loop length, is a critical determinant of transcription-translation coupling. While structural biology studies have indicated the existence of multiple expressomes with varying mRNA loop lengths, their *in vivo* roles and functional significance remain largely unexplored. This study delves into the mechanisms governing transcription termination within the *Escherichia coli* galactose operon, revealing a crucial role in the transcription and translation coupling state. The operon employs both Rho-independent and Rho-dependent terminators. Our findings demonstrate that long-loop coupled transcription-translation complexes preferentially terminate at the upstream Rho-independent terminator. In contrast, short-loop coupled complexes bypass the Rho-independent terminator and terminate at the downstream Rho-dependent terminator. The efficiency of the Rho-independent terminator is enhanced by an extended U-track, suggesting a novel mechanism for overcoming ribosome inhibition. These results challenge the traditional view of transcription termination as a random process, highlighting a predetermined mechanism contingent on the coupling state. This study emphasizes the intricate interactions between transcription and translation in prokaryotes. Understanding how these processes affect the RNA polymerase’s selection of transcriptional terminators is critical for developing strategies to regulate gene expression.

## 1. Introduction

In *Escherichia coli*, transcription and translation are coupled processes, with leading ribosome following behind RNA polymerase (RNAP) during protein synthesis (Burmann et al. 2010; Proshkin et al. 2010; Kohler et al. 2017; Wang et al. 2020; Blaha and Wade 2022; Woodgate and Zenkin 2023; Woodgate et al. 2024). This phenomenon, known as “coupling” of transcription and translation (CTT), is essential for regulating gene expression. Notably, CTT is a dynamic process rather than a static one. Structural studies have significantly advanced our understanding of the molecular basis of CTT by providing valuable insights (Kohler et al. 2017; Wang et al. 2020; Webster et al. 2020; Blaha and Wade 2022). The complex formed by the leading ribosome and the transcribing RNAP is known as the “expressome.” Within the expressome, the section of mRNA between the ribosome and the RNAP activity center is referred to as the mRNA loop. Expressome can be classified into factor-assisted and factor-independent categories (Woodgate and Zenkin 2023). Factor-assisted expressome involves the participation of NusG and NusA, which restrict the relative movement of ribosomes and RNAP. The mRNA loops typically measure approximately 41-47 nucleotides in length (Wang et al. 2020). While longer mRNAs were not experimentally tested, the authors theorized that in factor-assisted expressomes, the machinery could potentially accommodate up to 50 nucleotides of mRNA between their active centers. Conversely, factor-independent expressomes lack NusA and NusG, relying solely on direct ribosome-RNAP interactions. These complexes exhibit closer proximity, with the ribosome positioned approximately 26-34 nucleotides from the RNAP active center (Kohler et al. 2017; Wang et al. 2020; Webster et al. 2020). Structural studies of these “collided” complexes indicate incompatibility with NusG/NusA binding and reduced capacity for mRNA looping.

RNAP initiates RNA synthesis at the promoter and terminates transcription, releasing the nascent RNA molecule. Accurate and precise regulation of transcription termination is essential for maintaining normal physiological functions across all organisms. Two primary mechanisms govern transcription termination: intrinsic termination, or Rho-independent termination (RIT) (Gusarov and Nudler 1999; Yarnell and Roberts 1999; Ray-Soni et al. 2016; Roberts 2019; Gupta and Pal 2021), and Rho-dependent termination (RDT) (Roberts 1969; De Crombrugghe et al. 1973; Ciampi 2006; Cardinale et al. 2008; Ray-Soni et al. 2016; Jain et al. 2019; Roberts 2019; Song et al. 2022). RIT relies on a transcriptional pause induced by RNAP at the DNA’s RIT site (Gusarov and Nudler 1999; Epshtein et al. 2007; Weixlbaumer et al. 2013; Vvedenskaya et al. 2014). This pause allows the formation of a hairpin structure, the terminator hairpin, within the paused RNAP’s exit channel (Kang et al. 2018; Kang et al. 2019). The terminator hairpin stabilizes the transcriptional pause at the RIT site and weakens the U-A base pairing between the transcribed RNA and DNA template, disrupting the transcription complex. Ultimately, RNAP and the transcribed RNA dissociate from the DNA (Komissarova et al. 2002; Komissarova et al. 2008; Lubkowska et al. 2011; Kang et al. 2018; You et al. 2023). Transcriptional pausing of RNAP is also essential for RDT (Kassavetis and Chamberlin 1981; Wang et al. 1995; Weixlbaumer et al. 2013; Larson et al. 2014; Vvedenskaya et al. 2014). This pausing occurs downstream of the Rho utilization (*rut*) site, a cytosine-rich (C-rich) region (Richardson and Richardson 1996; Skordalakes and Berger 2003). The Rho termination factor, a protein, binds to this C-rich region within the transcribed RNA, a stretch of 73 nucleotides in the case of the galactose operon, in the absence of ribosomes (Alifano et al. 1991; Gocheva et al. 2015; Washburn and Gottesman 2015; Ray-Soni et al. 2016; Nadiras et al. 2018; Wang et al. 2019). The interaction between RNAP and Rho ultimately leads to the release of both RNAP and the transcribed RNA from the DNA template (Adhya and Gottesman 1978; Adhya et al. 1979; Adhya 2003; Epshtein et al. 2010; Zhu et al. 2019; Hao et al. 2021; Said et al. 2021; Molodtsov et al. 2023). It is well-established that translation significantly influences transcription termination (Zurawski et al. 1978; Roland et al. 1988; Wang et al. 2019; Jeon et al. 2021; Jeon et al. 2022). RIT is primarily determined by the distance between the translation stop codon and the transcript RNA’s terminator hairpin (Roland et al. 1988; Gusarov and Nudler 1999; Komissarova et al. 2008; Irnov and Winkler 2010; Li et al. 2016; Ray-Soni et al. 2016; Zhang and Landick 2016; Roberts 2019), while RDT occurs when transcription and translation become uncoupled (Adhya and Gottesman 1978; Zurawski et al. 1978; Roland et al. 1988; Wang et al. 2019; Jeon et al. 2021; Jeon et al. 2022).

Our previous research focused on regulating transcription termination within the *E. coli* galactose operon, comprising four structural genes: *galE, galT, galK*, and *galM*. The operon employs multiple internal Rho-dependent terminators located downstream of each gene’s open reading frame (ORF) (*galE, galT*, and *galK*) (De Crombrugghe et al. 1973; Wang et al. 2014a; Jeon et al. 2021; N et al. 2023; Wang et al. 2023). These RDTs act as safeguards to prevent “run-through” transcription when translation is disrupted. For example, a failure in *galT* translation initiation triggers transcriptional and translational decoupling, leading to RDT activation downstream of *galE* (Jeon et al. 2022). Similarly, *galK* translation failure activates sRNA Spot 42 binding, RDT downstream of *galT*, and subsequent mRNA degradation by RNase E (Moller et al. 2002; Wang et al. 2015; Jeon et al. 2020). At the operon’s terminus, the Rho-independent terminator and Rho-dependent terminator function concurrently. RIT occurs 30 nucleotides downstream of the *galM* translation stop codon, while RDT is positioned 94 nucleotides downstream of RIT. A portion of the transcript undergoes RIT-mediated termination, with the remainder terminating via RDT. Our earlier findings established the independence of RIT and RDT, with both processes regulated by NusA and NusG (Wang et al. 2019). Evidence from synthetic sRNA studies suggests varying translation densities within the galactose operon, with higher density in the *galET* region compared to the downstream *galKM* region (Wang et al. 2023), indicating that CTTs within the galactose operon may incorporate multiple mRNA loop length.

While structural biology studies have indicated the existence of multiple expressomes with varying mRNA loop lengths, their *in vivo* roles and functional significance remain largely unexplored. This study utilized the *E. coli* galactose operon as a model system to demonstrate the existence of diverse CTT complexes under physiological conditions. Our findings revealed that mRNA loop length significantly influences the choice between RIT and RDT. Specifically, CTTs with longer mRNA loops preferentially terminate at the RIT site, while those with shorter loops bypass RIT site and proceed to the downstream RDT site. These results suggest that the galactose operon’s Rho-independent terminator and Rho-dependent terminator have evolved to accommodate CTTs with varying mRNA loop lengths. The observed correlation between mRNA loop length and termination site selection suggests a potential regulatory mechanism that could influence gene expression at a global level. This mechanism could contribute to the fine-tuning of gene expression in response to various environmental and developmental conditions.

## 2. RESULTS

### 2.1. Experimental design to isolate and quantify RIT efficiency

To specifically investigate the efficiency of the RIT in the galactose operon, we introduced a hairpin structure downstream of the RIT site to block 3’ to 5’ exonuclease digestion (Jeon et al. 2022). This was necessary because, in the wild-type *E. coli* strain MG1655, transcriptional termination downstream of the *galM* gene occurs at both the RIT and the RDT sites, followed by 3’ to 5’ exonuclease digestion, resulting in a stable 4,313 mRNA 3’ end (Wang et al. 2019). By blocking exonuclease digestion, we could isolate the effects of the RIT. This strategy was implemented by constructing plasmid pRITHP, which contains the entire galactose operon, the RIT coding sequence, and the downstream hairpin coding sequence (Figure 1A). We then mutated the G and C nucleotides at the bottom of the Rho-independent terminator stem to C and G in pRITHP, disrupting the Rho-independent terminator hairpin structure and creating the Rho-independent terminator-inactivated mutant plasmid pRIT°HP. As a control, we also constructed plasmid pWT, which contains the intact galactose operon with both the Rho-independent terminator and the Rho-dependent terminator intact (Figure 1A). These plasmids were individually transformed into a Δ*gal* strain lacking the chromosomal galactose operon to create Δ*gal*-pWT, Δ*gal*-pRITHP, and Δ*gal*-pRIT°HP strains for subsequent analysis.

**Figure 1.**
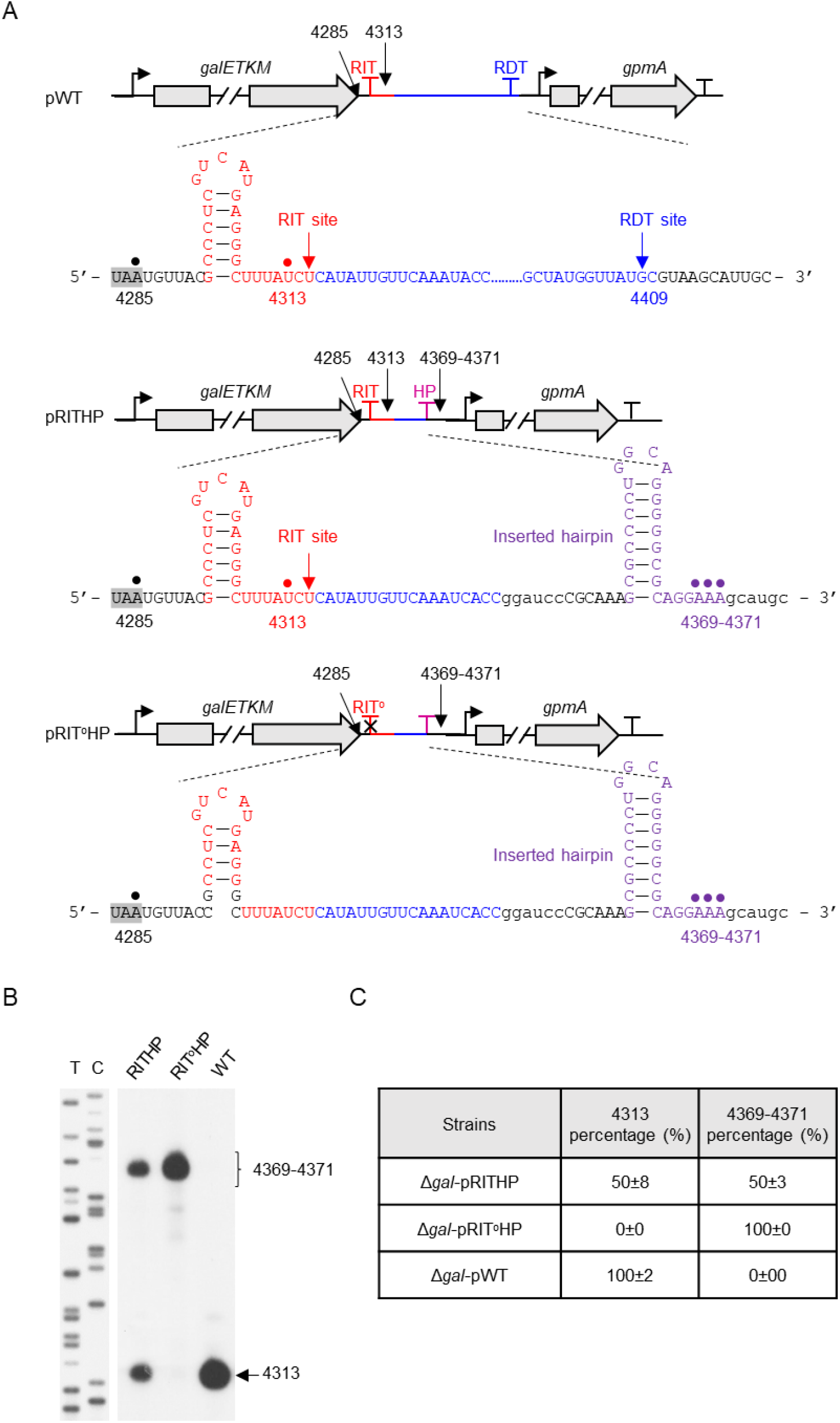
Transcription termination and passaging assay of RIT. (A) Schematic representation of pWT, pRITHP, and pRIT°HP constructs. The RIT signal (terminator hairpin) is displayed as a red line and T arrow, and the RDT signal (*rut* site coding sequence) is shown as a blue line and T arrow. The RIT signal of *gpmA* is presented as a black T arrow. Inserted hairpins are denoted by a purple T arrow in pRITHP and pRIT°HP. Mutations in the two G: C base pairs at the bottom of the hairpin stem were mutated and RIT inhibition was shown by a black cross. The translation terminator UAA of the *galM* gene is highlighted with a gray background. Additional sequences that were incorporated during the construction of the plasmid are represented in lowercase letters. (B) Visualization of the 3’ ends by 3’ RACE assay and primer extension. The 3’ end of 4313 shows the amount of RIT-terminated transcript, while the 3’ ends 4369-4371 show the amount of transcription passing through RIT. T/C: DNA sequencing ladders. (C) Percentage of 4313 and 4369-4371 3’ ends based on calculations in (B). The signal intensity of the bands was quantified using ImageJ software.

To assess RIT efficiency, we employed 3’ RACE-PE (rapid amplification of cDNA 3’ ends and primer extension) to quantify the abundance of 4,313 3’ ends in the different strains (Lee et al. 2008; X et al. 2018). A 50% reduction in 4,313 3’ end formation was observed in Δ*gal*-pRITHP compared to Δ*gal*-pWT (Figure 1B, C), indicating that the Rho-independent terminator terminates approximately half of the transcription complexes. Concurrently, we detected the formation of 4,369-4,371 3’ ends, located downstream of the inserted hairpin in Δ*gal*-pRITHP (Figure 1B). This suggests that the inserted hairpin blocked 3’ to 5’ exoribonuclease digestion, allowing for the accumulation of these new 3’ ends. Notably, the combined levels of 4,313 and 4,369-4,371 3’ ends in Δ*gal*-pRITHP equaled those in Δ*gal*-pWT (Figure 1C), further supporting the notion that Rho-independent terminator terminates half of the transcription complexes. In the Δ*gal*-pRIT°HP strain, lacking a functional RIT, 4,313 3’ ends were absent, while 4,369-4,371 3’ end levels doubled compared to Δ*gal*-pRITHP (Figure 1B, C). These results confirm that RIT is responsible for the generation of 4,313 3’ ends and that half of the transcription complexes bypass the RIT site in the presence of a functional Rho-independent terminator.

### 2.2. The role of mRNA loop length in RIT efficiency

The galactose operon exhibits a 30-nucleotide distance between the *galM* translation termination codon and the RIT site’s 3’ end (Figure 2A). Given the distance between the RNAP surface and its active center, as well as the distance between the ribosome surface and its decoding center, which is approximately 12 nucleotides in each case (Kohler et al. 2017; You et al. 2023), the leading ribosome, when occupying the *galM* translation termination codon, physically hinders the formation of Rho-independent terminator hairpins. Consequently, under these circumstances, the leading ribosome effectively represses RIT.

**Figure 2.**
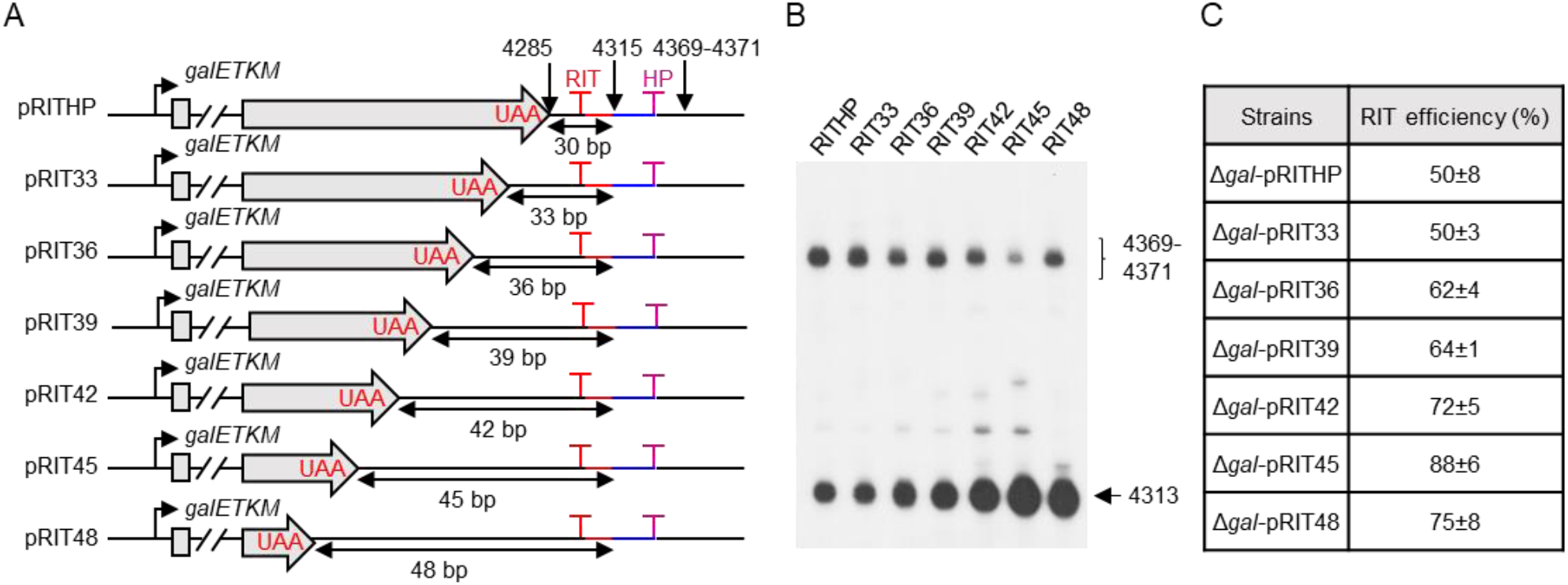
Assay of transcription terminated by and through Rho-independent terminator in a series of translation stop codon mutants. (A) Schematic presentation of pRITHP, pRIT33, pRIT36, pRIT39, pRIT42, pRIT45, and pRIT48 constructs. The RIT signal (the terminator hairpin) is displayed as a red line and T arrow, and the inserted hairpin is denoted by a purple T. The mutated stop codons (UAA) are shown in red and the distance between the translation stop codon of *galM* and the RIT site is shown by the double arrow. (B) Visualization of 3’ ends by 3’ RACE-PE assay. The 3’ end of 4313 shows the amount of transcript terminated by RIT, and the 3’ end of 4396-4371 represents the amount of transcription passing through RIT. (C) Relative RIT efficiencies are based on calculations in (B). The signal intensity of the bands was quantified using ImageJ software.

To investigate the relationship between mRNA loop length and RIT efficiency, we extended the distance between the *galM* translation stop codon and the RIT site from 30 to 48 nucleotides by moving the translation stop codon upstream while simultaneously mutating the original translation stop codon UAA to the lysine codon AAA in Δ*gal-*pRITHP (Figure 2A, Figure S1). 3’ RACE-PE analysis revealed a gradual increase in 4,313 3’ ends and a slight decrease in 4,369-4,371 3’ ends with the introduction of additional nucleotides (Figure 2B). Concurrently, RIT efficiency increased from 62% in Δ*gal*-pRIT36 to 88% in Δ*gal*-pRIT45, before declining to 75% in Δ*gal*-pRIT48 (Figure 2C). These findings suggest an optimal mRNA loop length of approximately 45 nucleotides for maximal RIT efficiency. We propose classifying mRNA loops shorter than 45 nucleotides as short-loop coupling, which inhibits RIT, and those longer than or equal to 45 nucleotides as long-loop coupling, which permits RIT. While the precise reasons for the efficiency decline beyond 45 nucleotides remain to be elucidated, these results highlight the influence of mRNA loop length on RIT.

### 2.3 Long-loop coupling complexes preferentially terminate at the Rho-independent terminator

We observed that in the Δ*gal-*pRITHP strain, approximately half of the transcripts terminated at the Rho-independent terminator despite its proximity (only 30 nucleotides distant) (Figure 1B, Figure 2B). This suggests that in approximately half of the CTTs, the leading ribosome is positioned sufficiently far from the Rho-independent terminator to allow hairpin formation. Two potential explanations for this phenomenon exist: either transcription and translation are uncoupled, or the mRNA loop within the coupled complex is equal to or longer than 45 nucleotides to allow the hairpin formation, that is, the long-loop coupling.

To differentiate between uncoupled and long-loop coupling complexes, we constructed a p*galM*-SD mutant by mutating the Shine-Dalgarno (SD) sequence of *galM* from GGAG to CCTC, effectively uncoupling transcription and translation (Jeon et al. 2020). Northern blot analysis revealed that the strain Δ*gal*-p*galM*-SD produced a significantly reduced mM1 band compared to the strain Δ*gal*-pWT (Figure 3A). To further elucidate the reasons for the decreased expression of the mM1 bands, we transformed GW20 (*ams1*^ts^), an RNase E temperature-sensitive strain, with pWT and p*galM*-SD plasmids and cultured them under both active (30°C) and inactive (44°C) RNase E conditions. Northern blot analysis demonstrated that while RNase E cleavage reduced mK1 and mM1 expression, mM1 transcript levels were significantly lower in the GW20-p*galM*-SD strain compared to the GW20-pWT strain, regardless of RNase E activity (Figure 3B). These results indicate that uncoupled transcription is inefficient in reaching the *galM* Rho-independent terminator, independent of RNase E activity. Consequently, we conclude that transcription and translation occur in a long-loop coupled state when RNAP encounters the *galM* Rho-independent terminator.

**Figure 3.**
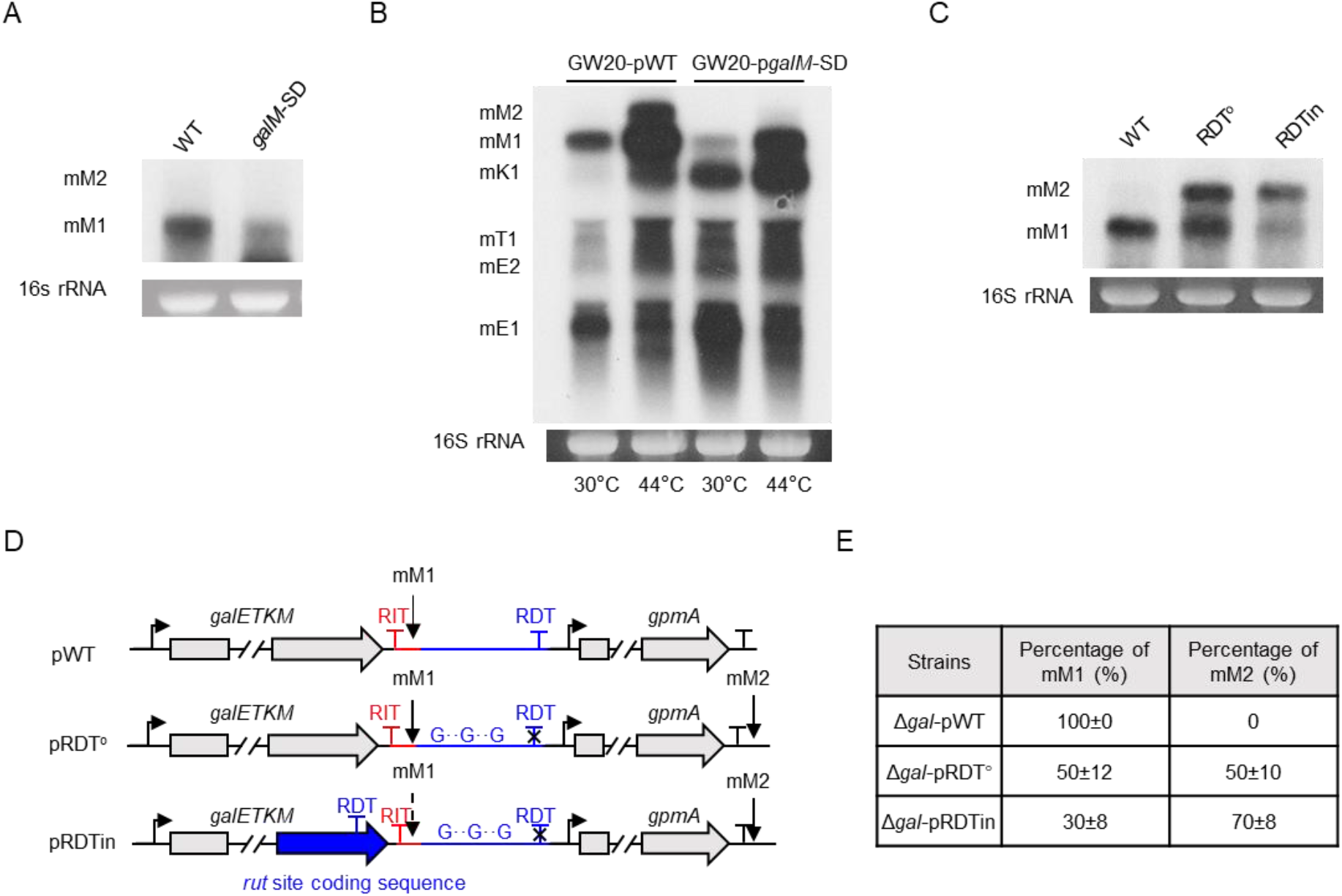
Long-loop CTTs terminate at the first terminator. (A) Northern analysis of full-length transcripts from Δ*gal-*pWT, and Δ*gal-*p*galM*-SD. (B) Northern analysis of galactose transcripts from GW20*-*pWT and GW20*-*p*galM*-SD under both active (30°C) and inactive (44°C) RNase E conditions. (C) Northern analysis of full-length transcripts from Δ*gal-*pWT, Δ*gal-*pRDT°, and Δ*gal-*pRDTin. (D) Schematic representation of pWT, pRDT°, and pRDTin constructs. The pRDT° plasmid was generated by replacing all 23 cytosines in the *rut* site coding sequence with guanine (G) to inhibit RDT. In pRDTin, the *rut* site coding sequence in pWT was inserted in-frame at the end of *galM*, upstream of the stop codon in pRDT°. The RIT signal (terminator hairpin) is displayed as a red T arrow. RDT signal is shown as a blue T arrow and its inhibition is shown by a black cross. (E) Percentage of mM1 transcripts calculated based on (C). The signal intensity of the bands was quantified using ImageJ software.

To reveal the existence of long-loop CTTs, we introduced a Rho-dependent terminator upstream of the *galM* Rho-independent terminator. We hypothesized that if long-loop CTTs exist, they would terminate at the inserted Rho-dependent terminator. To achieve this, we generated plasmid pRDT° by replacing all cytosines within the Rho-dependent terminator coding region of pWT with guanines (Wang et al. 2019). Subsequently, we constructed plasmid pRDTin by replacing a 132 nucleotide region upstream of the *galM* translation stop codon in pRDT° with the *galM* Rho-dependent terminator coding sequence (Figure 3D, Figure S2). Northern blot analysis revealed that Δ*gal*-pWT produced solely mM1 transcripts, while Δ*gal*-pRDT° generated equal amounts of mM1 and mM2 transcripts (Figure 3C). The absence of mM2 in Δ*gal*-pWT is attributed to the presence of the downstream RDT, which prevents transcription from reaching the *gpmA* gene (Wang et al. 2019). In contrast, introducing Rho-dependent terminator upstream of *galM* in the Δ*gal*-pRDTin strain markedly decreased mM1 transcript levels to 30% while concurrently inducing the production of mM2 transcripts (Figure 3C, E). We conclude that the inserted Rho-dependent terminator terminates a subset of long-loop CTTs, preventing their progression to the Rho-independent terminator. The remaining short-loop CTTs, unable to terminate at the RIT site, are processed to generate mM2 transcripts. These results corroborate the presence of long-loop CTTs that extend to the terminus of the galactose operon.

### 2.4 Short-loop CTTs inhibit RIT and terminate at the Rho-dependent terminator

To further investigate the termination behavior of short-loop CTTs, we examined the potential influence of the Rho-dependent terminator on transcription. Given that Rho-dependent terminators are located beyond the inhibitory range of the leading ribosome, we hypothesized that short-loop CTTs would undergo translation termination at the *galM* stop codon, followed by ribosome dissociation. Subsequently, RNAP would encounter and terminate at the downstream Rho-dependent terminator. In WT, transcripts terminated by the Rho-dependent terminator were rapidly degraded due to a lack of secondary structure, making it difficult for us to observe the initial RDT-terminated transcript. Therefore, we replaced the Rho-dependent terminator in pWT with a second *galM* Rho-independent terminator (RIT2), located 31 nucleotides downstream of the original RIT, generating plasmid pRIT2 (Figure 4A, Figure S3). This arrangement positioned the first Rho-independent terminator within the potential inhibitory range of the leading ribosome, while the second Rho-independent terminator was beyond this distance. Northern blot analysis of Δ*gal*-pRIT2 revealed a single predominant band (labeled mM1 or mM1* in Figure 4B), suggesting that a second Rho-independent terminator terminated RNAP transcription in short-loop CTTs. While theoretically, two transcripts terminating at the first and second RITs, respectively, should be present, their 30 bp length difference was indistinguishable due to the limitations of Northern blot resolution.

**Figure 4.**
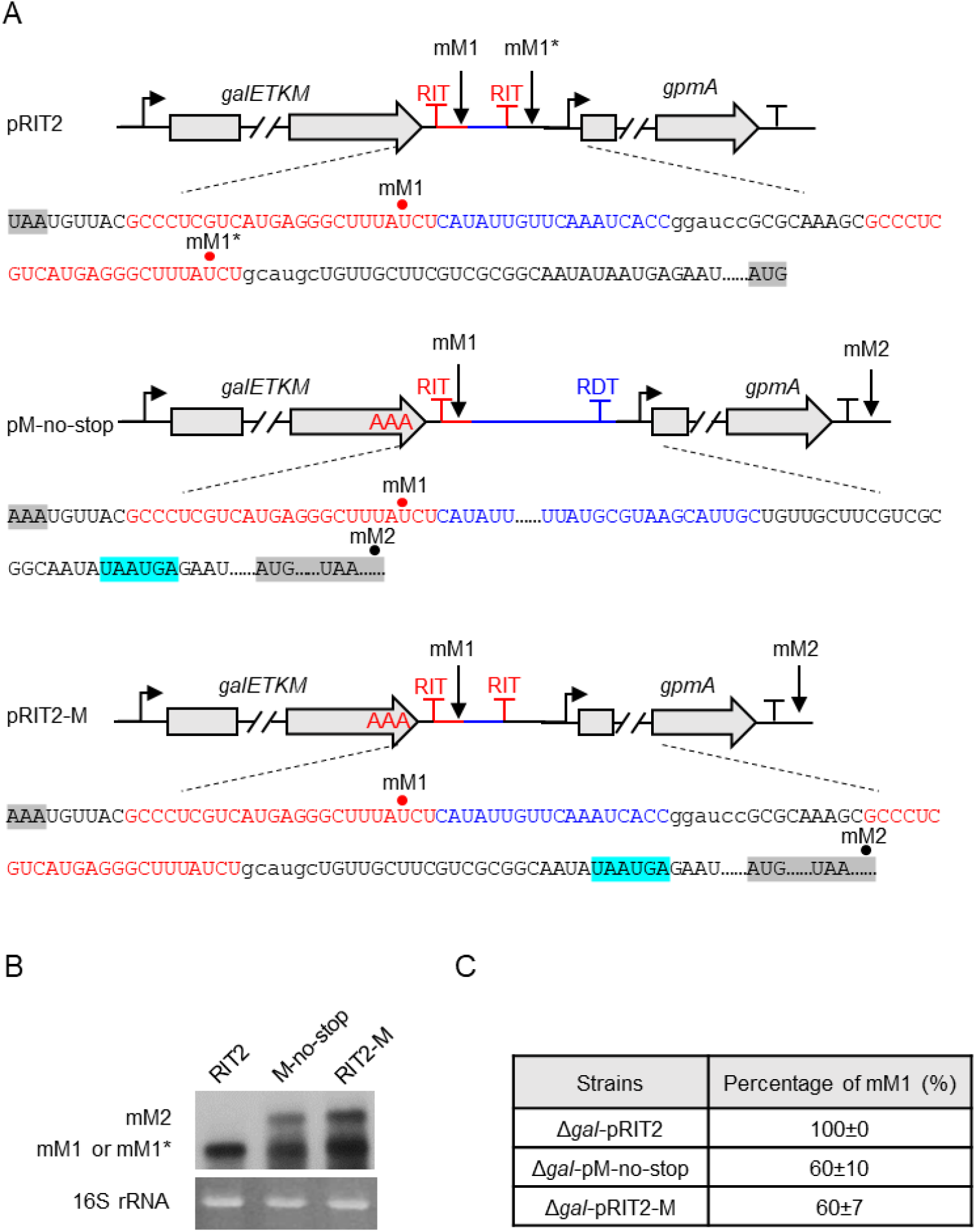
Short-loop CTTs terminate at the second terminator. (A) Schematic representation of pRIT2, pM-no-stop, and pRIT2-M constructs. The pRIT2 plasmid was generated by replacing the RDT coding sequence with the *galM* Rho-independent terminator coding sequence. In pM-no-stop, the translation stop codon of *galM* in pWT was mutated to AAA. In pRIT2-M, the translation stop codon *galM* in pRIT2 was mutated to AAA. The translation termination codon of the *galM* gene, as well as the translation start and termination codons of the *gpmA* gene, are displayed on a gray background. The two consecutive translation termination codons are shown on a blue background. (B) Northern analysis of full-length transcripts from Δ*gal*-pRIT2, Δ*gal*-pM-no-stop, and Δ*gal*-pRIT2-M strains. (C) Percentage of mM1 transcripts calculated based on (B). The signal intensity of the bands was quantified using ImageJ software.

We then mutated the *galM* translation termination codon UAA to lysine (AAA) in pWT and pRIT2 to generate the plasmids pM-no-stop and pRIT2-M, respectively (Figure 4A). These constructs permitted continued translation beyond the *galM* ORF. However, in both plasmids, we have identified two consecutive translation terminators with the sequence UAAUGA, located 154 base pairs downstream of the original translation terminator of the *galM* gene (Figure 4A). These terminators are situated in the frame of the GalM protein. We anticipate that this extended ORF will lead to translation termination at the newly discovered site. Considering that the new translation terminators are positioned downstream of the Rho-dependent terminator in pM-no-stop and the second Rho-independent terminator in pRIT2-M, we hypothesize that when RNAP encounters these terminators, transcription and translation will maintain a short-loop CTT state. This state is expected to inhibit the function of the second transcription terminator. Consequently, transcription is likely to extend beyond the *gpmA* gene, potentially resulting in the formation of an mM2 band.

As anticipated, Northern blot analysis of both strains demonstrated the presence of both mM1 and mM2 bands (Figure 4B). This finding indicates that the first terminator retains its ability to terminate long-loop CTTs, while the second terminator is less efficient. This inefficiency could potentially be due to the inhibitory effects of continued translation within the short-loop CTTs. Furthermore, in the Δ*gal*-pM-no-stop and Δ*gal*-pRIT2-M strains, the proportion of mM1 transcripts increased from 50% to 60% relative to the Δ*gal*-pRDT° strain (Figure 4C). This elevation in mM1 percentage could be attributed to transcription-translation uncoupling within the *gpmA* gene, triggered by the translation termination at UAAUGA. Consequently, this uncoupling leads to a diminished amount of RNAP reaching the downstream regions of the *gpmA* gene. Taken together, our findings support the existence of distinct populations of short- and long-loop CTTs. The termination behavior of these populations appears to be influenced by the coupling state and the presence of downstream terminators.

### 2.5 A terminator hairpin with eight uridines terminated short-loop CTTs

In a few operons, there were cases where the distance between the stop codon and the RIT site was less than 12 nucleotides, yet the occurrence of transcription termination was still observed (Johnson et al. 2020). We hypothesized that there are three possibilities: first, the Rho-independent terminator can overcome the inhibitory effect of the leading ribosome; second, the transcript terminated by the downstream Rho-dependent terminator is processed by the exoribonuclease near the hairpin structure, resulting in the “artifact” that the Rho-independent terminator played a transcription termination role. In the third case, the Rho-independent terminator acts not only as a transcription terminator but also as a hairpin structure that maintains RNA stability. The latter two cases have been confirmed in our previous work (Wang et al. 2019) and Dar et al. (Dar and Sorek 2018), and here we would like to explore whether the first scenario holds.

To investigate whether a Rho-independent terminator with a longer U-track could overcome the inhibitory effects of the leading ribosome on RIT, we mutated the U-track of the *gal* Rho-independent terminator in pWT and pRDT° to contain eight uridines, generating p8Us and pRDT°8Us plasmids, respectively (Figure 5A, B). Northern blot analysis revealed that only mM1 transcripts were detected in both Δ*gal*-pWT and Δ*gal*-p8Us strains. However, while the Δ*gal*-RDT° strain typically produced mM1 transcripts in approximately 50% of CTTs, the Δ*gal*-pRDT°8Us strain exhibited mM1 transcripts in nearly 90% of CTTs (Figure 5C, D). These results suggest that extending the U-track in the Rho-independent terminator increases its efficiency in terminating transcription, even in the presence of a ribosome stalled at the stop codon. This enhanced termination likely occurs due to a prolonged RNAP pause induced by the extended U-track, providing sufficient time for ribosome release and subsequent formation of the termination stem-loop.

**Figure 5.**
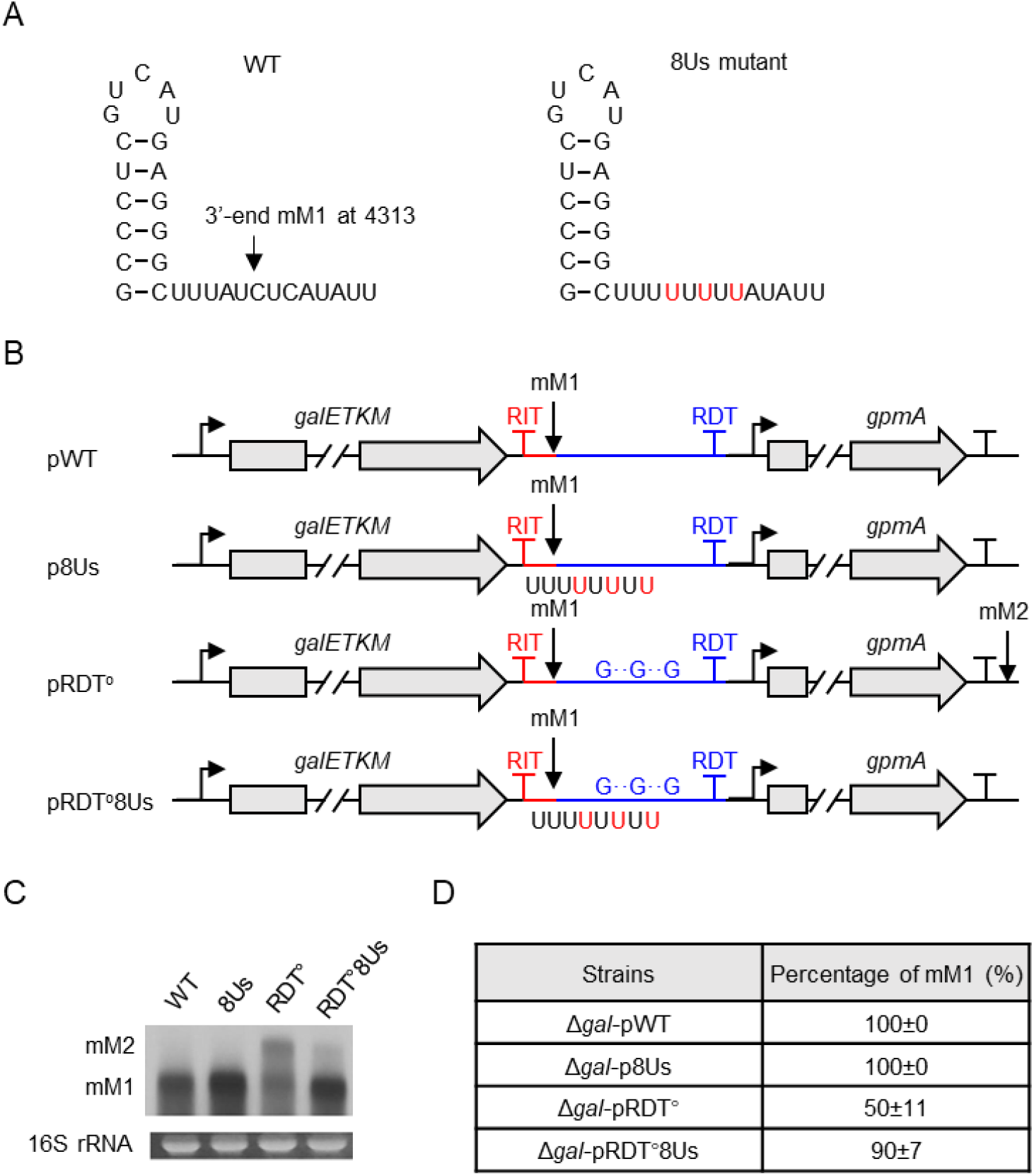
Termination of short-loop CTTs by the *galM* terminator hairpin with 8Us. (A) Schematic representation of the terminator hairpin at the terminus of *galM* and a mutant terminator hairpin with eight consecutive uridines (8Us) from the hairpin stem. (B) Schematic representation of pWT, p8Us, pRDT°, and pRDT°8Us constructs. The p8Us plasmid was generated by replacing WT Rho-independent terminator in pWT with 8Us, and the pRDT°8Us plasmid was generated by replacing WT Rho-independent terminator in pRDT° with 8Us. (C) Northern analysis of full-length transcripts of the galactose operons of Δ*gal-*pWT and Δ*gal-*pRDT°, and corresponding mutants that harbor the terminator hairpin with 8 Us, Δ*gal-*p8Us and Δ*gal-*pRDT°8Us. (D) Percentage of mM1 transcripts calculated based on (B). The signal intensity of the bands was quantified using ImageJ software.

## 3. DISCUSSION

This study uncovers the sophisticated mechanisms that govern the transcriptional termination of the galactose operon. By examining different mRNA loop lengths—short-loops (less than 45 nucleotides) and long-loops (45 nucleotides or more)—we reveal the operon’s adaptability in response to these variations. This categorization is grounded in structural biology insights: the distance from the RNAP surface to its active center is approximately 12 nucleotides (You et al. 2023), similar to the distance from the ribosome surface to its decoding center (Kohler et al. 2017). Moreover, the *galM* Rho-independent terminator hairpin in the galactose operon spans 17 nucleotides, suggesting that a minimum loop length of about 41 nucleotides is necessary for Rho-independent terminator RNA hairpin formation. Our experimental results corroborate this structural prediction, indicating that the peak efficiency of RIT occurs at an mRNA loop length of 45 nucleotides. This alignment between experimental data and structural biology validates our approach and underscores the importance of mRNA loop length in RIT efficiency. Additionally, we propose that the lengths of short- (inhibitory termination) and long- (permissive termination) loops may vary across different transcription terminators due to differences in stem-loop structures. This variability suggests a potential mechanism for fine-tuning gene expression through alterations in stem-loop lengths.

Based on these findings, we propose a molecular model for transcription termination at the galactose operon’s terminus. Two types of CTTs reach the terminus: In one case, long-Loop CTTs, approximately 50% of these complexes allow the terminator hairpin to form in the RNAP exit channel, causing the RNAP to pause and terminate at RIT. Here, transcription may terminate before translation does (Figure 6A). In the second case, short-loop CTTs, about 38% of the complexes exhibit coordinated yet independent movement of RNAP and the leading ribosome. The ribosome reaches the translation stop codon before the Rho-independent terminator hairpin forms, inhibiting hairpin formation and preventing RIT. Transcription resumes downstream, with translation likely terminating before transcription does, leading to decoupling and potential Rho-dependent termination (Figure 6B). Interestingly, a subset of transcription bypasses the RIT site regardless of the distance between the stop codon and the RIT site, suggesting that Rho-independent terminators are not 100% efficient. This is reflected in the inability of the transcription complex to terminate at the *galM* operon’s terminus.

**Figure 6.**
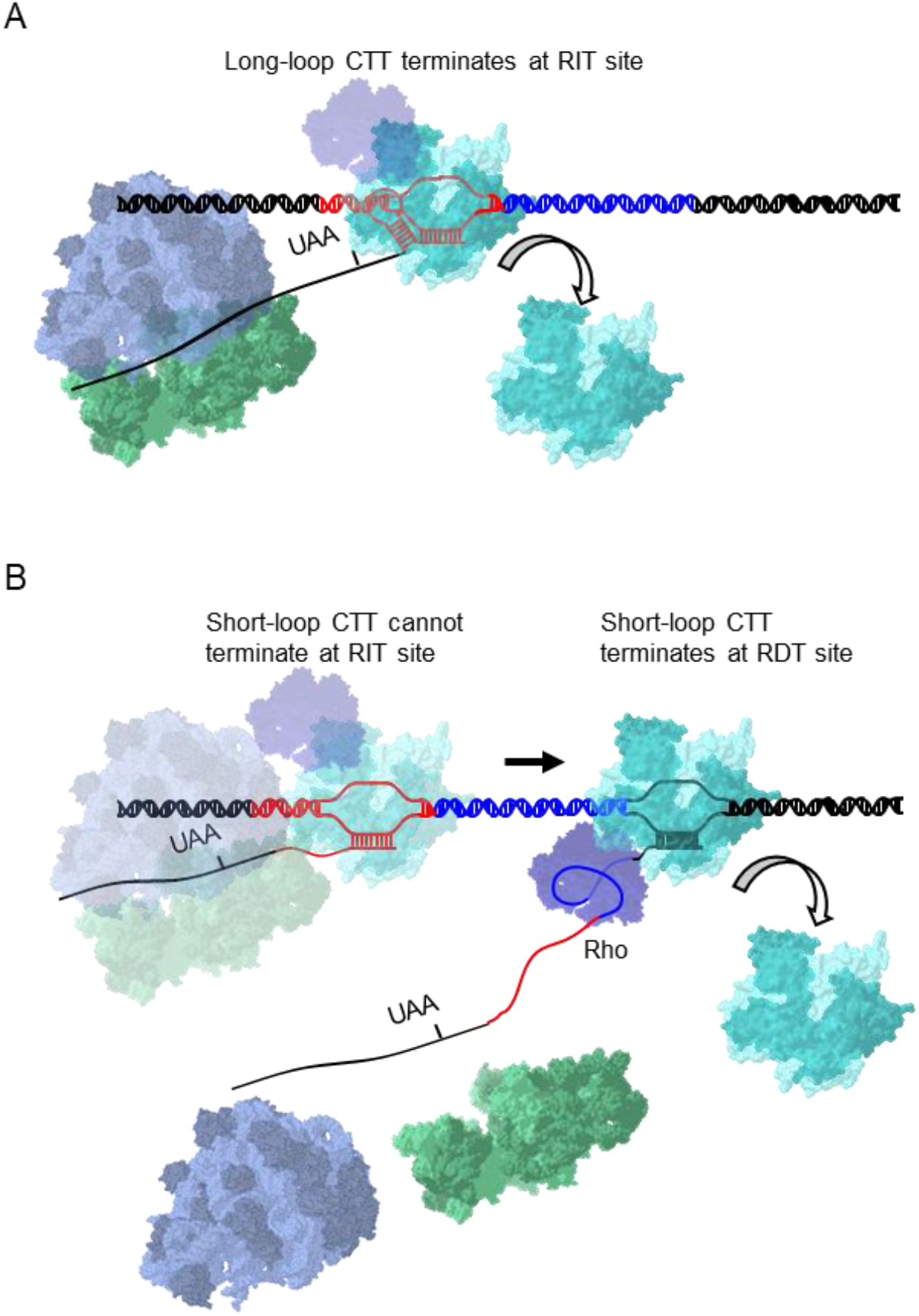
Termination Model. (A) Long-loop CTT: In this scenario, the distance between the decoding center of the leading ribosome (purple, the 50S subunit) and green (the 30S subunit) and the active center of RNA polymerase (cyan structure) is longer than the distance between the stop codon of *galM* (denoted as UAA) and the 3’ end of the transcript RNA at the active site of the RNA polymerase. The terminator hairpin coding sequence is shown in red and the *rut* site coding sequence is shown in blue. The terminator hairpin on mRNA is also shown in red. The leading ribosome of this structure is not in a position to inhibit the formation of a terminator hairpin at the exit channel of the RNA polymerase. Therefore, the terminator hairpin forms and RIT occurs. (B) Short-loop CTT: In this scenario, the leading ribosome inhibits the formation of a terminator hairpin. Therefore, RNA polymerase can resume transcription and RIT does not occur. Rho (violet structure) is pre-bound to RNAP and contacts *rut* once it emerges from the RNAP exit channel and terminates transcription.

In conclusion, our model suggests that the galactose operon’s Rho-independent and Rho-dependent terminators have adapted to accommodate CTTs with varying coupling statuses. The Rho-independent terminator is designed for long-loop coupled terminators, while the Rho-dependent terminator handles short-loop coupled ones. Our findings point towards the significance of examining the sophisticated molecular interactions at play between ribosome and RNAP. We propose that future studies should delve deeper into these interactions to gain more insight into the dynamic processes of transcription and translation. By gaining insights into the dynamic processes that govern mRNA loop length regulation, we can manipulate gene expression in more precise ways. The enhanced understanding afforded by this finding will not only refine our comprehension of cellular processes but also hold the promise of illuminating new aspects of molecular biology.

## 4. EXPERIMENTAL PROCEDURES

### 4.1 Bacterial strains and growth conditions

The MG1655 Δ*galETKM*, GW20 (*ams1*^ts^), an RNase E temperature-sensitive strain and EPI300 strains were used in this study. Chromosomal deletion strains of the corresponding MG1655 genes were created using red-mediated recombination (Datsenko and Wanner 2000). The primers and their respective sequences used are listed in Table S1. Cells harboring the described plasmids were grown at 37 °C in Lysogeny Broth (LB) medium supplemented with 0.5% (wt/vol) galactose and chloramphenicol (20 µg/mL).

### 4.2 Plasmid cloning

The pHL1703, p*gal-gpmA* (pWT, in this study) plasmid was made by insertion of the galactose operon and the downstream gene, *gpmA* (from-75 to+5,397), between the *Eco*RI and *Bam*HI sites of pCC1BAC (Epicentre Biotechnologies) (Wang et al. 2019). The plasmids used in the study are shown in Table S2. The DNA fragment was obtained via PCR amplification of the genomic DNA with the corresponding primer pairs as listed in Table S1. The mutagenic DNA fragments were then obtained using PCR amplification of the pWT, and employed as a “mega primer” for the next round of PCR. The resulting PCR fragments were digested with *Mlu*I and *Bam*HI and then ligated into pHL1277 (Wang et al. 2014b) to generate pRDT°, p8Us, and pRDT°8Us. To construct the pRDTin plasmid, a pHL1933 plasmid containing the galactose operon sequence from *gal* coordinates -73 to +4,144 was first created. The DNA fragment encoding the *rut* site coding sequence (138 bp) was introduced between the *BamH*I and *Sph*I sites of pHL1933. The DNA fragments containing the *gpmA* sequence and the “*rut* site coding region mutations” fragment were simultaneously inserted into the *Sph*I site.

For the construction of pRITHP, the DNA fragment of the *galE* hairpin (Jeon et al. 2022) was ligated between *Bam*HI and *Sph*I of pHL1277 (Wang et al. 2014b). For localized mutagenicity, synthetic primers containing the desired mutations have been designed. After obtaining the mutagenic DNA fragments through PCR amplification of the pWT plasmid, the DNA fragments were used as a “mega primer” for the next round of PCR (Tyagi et al. 2004). The resulting PCR fragments were digested with *Mlu*I and *Bam*HI and then ligated into pRITHP to generate pRIT°HP, pRIT33, pRIT36, pRIT39, pRIT42, pRIT45, and pRIT48.

### 4.3 RNA preparation

Direct-zol™ RNA MiniPrep (Zymo Research) kit was used to purify the total RNA from clarified cell lysates. Cell lysates were generated by harvesting equal numbers of cells (2×10^8^) at an OD_600_ of 0.6. The RNA was subsequently isolated as recommended by the manufacturer. RNA was eluted in 30 μL of RNA storage buffer (Thermo Fisher Scientific) and quantified by measuring the absorbance at 260 nm using a NanoDrop (Thermo Fisher Scientific) spectrophotometer.

### 4.4 3’ RACE-PE assay

For the 3’ RACE assay, RNA ligation was performed at 37 °C for 3 h in a 25 μL reaction volume containing 2.5 µg of total RNA, 100 nM synthetic RNA oligomer possessing a 5’-phosphate and 3’-inverted deoxythymidine (27 nucleotides; Dharmacon), 5 U of T4 RNA ligase (Thermo Fisher Scientific), and 10 U of rRNasin (Promega). One microgram of RNA (eluted from the G-50 column (GE Healthcare)) was reverse transcribed at 37 °C for 2 h in a 20 μL reaction volume containing 4 U of Omniscript reverse transcriptase (Qiagen), 0.5 mM each dNTP, 0.4 M 3RP primer complementary to the RNA oligomer (Table S1), and 10 U of rRNasin.

A 2 μL of the cDNA was used as the template for PCR amplification with gene-specific primers and the 3RP primer (Table S1) using HotStar Taq DNA polymerase (Qiagen). To assay the 3’ ends of the *gal* mRNAs, the amplified cDNA was purified and used as a template for a primer extension reaction (X et al. 2018). The primer extension reaction was performed in a volume of 20 μL with a ^32^P-labeled primer (complementary to *galM* of the galactose operon mRNA; Table S1) and 1 U of Taq polymerase (Qiagen) (X et al. 2018; N and Lim 2022). The reaction products were resolved on an 8% sequencing gel for 2 h at 60 W, and the radioactive bands were visualized after exposure to X-ray film (Duksan (DS) Lab).

### 4.5 Northern blot assay

Total RNA (10 μg with 10 μg/ml ethidium bromide) was resolved by 1.2% (wt/vol) formaldehyde-agarose gel electrophoresis at 8 V/cm for 2 h. RNA integrity was assessed under UV light after electrophoresis. Then, RNA was transferred overnight to a positively charged nylon membrane (Thermo Fisher Scientific) using a downward transfer system (TurboBlotter; Whatman). The RNA was then fixed to a nylon membrane by baking at 80 °C for 1 h. The blot was pre-hybridized in 7 mL ULTRAhyb® Hybridization Buffer (Thermo Fisher Scientific) at 65° C for 30 min. The ^32^P-labeled DNA probe (See below) was denatured at 95 °C for 5 min, and then 5 μL of the probe was added to the hybridization buffer. The hybridization was performed at 42 °C overnight. Then, the blot was washed twice in low-stringency wash buffer (2xSSC, 0.1% SDS) for 5 min each at room temperature and then twice in high-stringency wash buffer (0.2xSSC, 0.1% SDS) for 15 min each at 42 °C. Finally, the radioactive bands were visualized after exposure to X-ray film.

### 4.6 Isotopic labeling of DNA probe

The Northern blot probe was prepared as follows. First, a 500-bp DNA fragment in the *galE* region (from +27 to +527) was prepared by PCR using primers indicated in Table S1. ^32^P-labeled DNA probes were produced as follows: Briefly, the template DNA (0.15 pmol) was mixed with random hexamer (4 μL of 1 mM) in a total volume of 28 μL and then heated at 95 °C for 3 min. Then, the reaction was rapidly cooled for 5 min on ice, and then 5 μL 10× Klenow fragment buffer (Takara), 5 μL dNTP mix (0.2 mM dATP, dGTP, and dTTP), 10 μL α ^32^P-dCTP, and 2 μL Klenow fragment (2 U/μL) was added to the mixture. The reaction was performed by incubation at 37 °C for 1 h, and then the Klenow fragment was inactivated at 65 °C for 5 min. The product was purified via passage through the G-50 column (GE Healthcare).

### 4.7 Quantification and Statistical Analysis

Each experiment was independently repeated at least three times with similar results. The signal intensities of the bands were quantified using the software ImageJ (Schneider et al. 2012).

## AUTHOR CONTRIBUTIONS

Conceptualization, funding acquisition, and supervision: X.W., H.J., and H.L.; Formal analysis, data curation, investigation, methodology, validation, visualization, and data analysis: M.N., and X.W.; Writing - original draft preparation: X.W., and H.L., Writing - review, and editing: M.N., X.W., and H.L.

## ACKNOWLEDGEMENTS

This work was supported by the National Natural Science Foundation of China (31971339 and 32171422), the National Key Research and Development Program of China (2022YFF1000700), and the Fundamental Research Funds for the Central Universities (2662022SKYJ004). This work was also supported by the research fund of Chungnam National University.

## DATA AVAILABILITY STATEMENT

All relevant data are within the paper and its supplemental file.

## ADDITIONAL FILES

### Supplementary Material

**Supplementary material** (Appendix, PDF file). Figures S1 to S3 and Tables S1 to S2

